# Colon resident CD8+ T cells from people with HIV on antiretroviral therapy exhibit mitochondrial dysfunction and impaired responses

**DOI:** 10.1101/2023.05.02.539176

**Authors:** Upasana Das Adhikari, Palak V. Shah, Leah M. Froehle, Alice H. Linder, Meaghan Flagg, Crystal Rawlings, Muntsa R. Junca, Lilian Folarin, Farina Karim, Jonathan Li, Henrik Kloverpris, Douglas S. Kwon

**Affiliations:** Ragon Institute of Massachusetts General Hospital, Massachusetts Institute of Technology, and Harvard, Cambridge, Massachusetts, USA; Harvard Medical School, Boston, Massachusetts, USA; Africa Health Research Institute (AHRI), Durban, South Africa; Department of Immunology and Microbiology, University of Copenhagen, Denmark; University College London, Division of Infection and Immunity, London, United Kingdom; Division of Infectious Diseases, Massachusetts General Hospital, Boston, Massachusetts, USA

## Abstract

CD8+ tissue-resident memory T cells (CD8+ TRM) play an important role in mediating immune responses against HIV in the gastrointestinal (GI) mucosa. Antiretroviral therapy (ART) successfully supresses HIV replication in the blood, but fails to restore GI homeostasis, particularly within gut associated lymphoid tissue (GALT). The importance of cellular metabolism in shaping CD8+ TRM antiviral function has emerged as a crucial aspect of general immunoregulation. To gain better insight into the immunoregulation of CD8+ TRM during HIV, we characterized the function of blood and colon CD8+ T cells in people with HIV (PWH) on ART and HIV-uninfected individuals. We show that a higher proportion of activated CD8+ TRM persisted in the colon of PWH on ART, and these cells exhibited weaker immune response and compromised mitochondrial function compared to their peripheral counterparts. We also observed decreased mitochondrial function in CD8+ T cell in the colon compared to peripheral blood in PWH on ART. The study indicates that mitochondrial bioenergetic properties may be an indicator of colonic CD8+ TRM dysfunction in PWH on ART and highlights the importance of understanding cellular metabolism in shaping CD8+ TRM function to develop targeted immunotherapies for PWH.

## INTRODUCTION

During HIV infection, CD8+ T cells play a crucial role in recognizing and killing infected cells to control viral replication and limit disease progression^1^. Tissue-resident memory CD8+ T cells (CD8+ TRM) reside in mucosal tissues such as in the gastrointestinal (GI) tract^2, 3^ and the failure of these cells to control viral replication in gut associated lymphoid tissue (GALT) leads to loss to intestinal homeostasis, which is a hallmark of HIV infection. This results in breakdown of intestinal barrier function, translocation of bacterial products into the systemic circulation, and chronic immune activation^4^. Treatment with antiretroviral therapy (ART) successfully suppresses viral replication in the periphery, but fails to reverse intestinal damage and associated systemic inflammation^5, 6^, and studies have shown that the GALT harbors ongoing viral replication despite successful long-term ART^7, 8^. Although recent studies have highlighted the critical role of CD8+ TRM in maintaining intestinal homeostasis and protecting against infections in the GI mucosa, the functional profile of these cells in people with HIV (PWH) on ART remains incompletely understood^9–11^.

Immunometabolism is a critical regulator of CD8+ T cell function, and mitochondrial properties such as mitochondrial mass, mitochondrial membrane potential, and spare respiratory capacity (SRC) control cellular activation and effector functions^12–17^. Despite long-term suppressive ART, CD8+ T cell responses in PWH remain dysfunctional^18^ with impairment associated with higher mitochondrial mass^19, 20^. To meet the bioenergetic demands of cellular activation, CD8+ T cells in the peripheral blood of PWH undergo metabolic reprogramming, switching from mitochondrial oxidative phosphorylation (OXPHOS) to aerobic glycolysis^21, 22^ indicating the importance of mitochondrial function in antiviral CD8+ T cell responses. The failure of metabolic shift to mitochondrial OXPHOS and the ability to utilize fatty acid oxidation in CD8+ T cells in the peripheral blood of PWH can abrogate the conversion to CD8+ T cell memory following peak effector differentiation and contraction^23^. Therefore, targeting mitochondria may be a promising strategy for improving immune function in PWH. However, most studies on mitochondrial function in PWH have focused on the characterization of circulating cells, leaving an incomplete understanding of the mitochondrial bioenergetic properties CD8+ TRM. Such studies are likely to reveal compartment-specific differences in CD8+ T cell mitochondrial function, providing a more detailed understanding of immune dysregulation in PWH.

We investigated mitochondrial metabolism and immune function of blood and colonic CD8+ T cells in PWH receiving ART. Despite effective control of HIV replication in the peripheral blood, we found elevated levels of CD8+ T cell activation in the colon of PWH on ART, which was associated with an increased marker of bacterial translocation. We also observed that CD8+ T cells from the colon exhibited increases in mitochondrial membrane potential and impaired immune effector function compared to T cells in the peripheral blood in PWH on ART. Transcriptional analysis of CD8+ T cells showed that the mitochondrial OXPHOS pathway was downregulated in the blood and colon of PWH on ART, with decreased expression of genes associated with the mitochondrial electron transport chain (ETC) between colon and peripheral blood CD8+ T cells. Additionally, mitochondrial respiration was significantly dampened in colonic CD8+ TRM compared to peripheral CD8+ T cells and those from HIV-uninfected donors. Overall, the study suggests that there is tissue-specific dysfunction of mitochondrial responses in CD8+ TRM in ART-treated PWH, and that these cells are more vulnerable to metabolic dysregulation during HIV infection.

## RESULTS

### Persistent hyperactivation of colon CD8+ TRM in PWH on ART

CD8+ TRM play a critical role in the immune response against HIV in GALT, but despite effective treatment with ART, there are persistent defects in intestinal homeostasis. To address this, we characterized a cohort of PWH on ART and HIV-uninfected individuals who underwent sampling of matched blood and colonic tissue obtained via lower endoscopy (**Table 1**). Our cohort was predominantly male with a median age of 50 years. There were no statistically significant differences between age, sex, self-identified race, or ethnicity between the PWW and HIV-uninfected individuals. PWH were all stably suppressed on ART with undetectable plasma viral loads and a median duration of treatment of approximately 7 years (**Table 1**). Participants were screened for any indications of GI or other inflammatory disease and those with significant clinical findings during endoscopy were excluded from further analysis.

Table 1.

Table 1
| Patient Characteristics | HIV Negative (n = 46) | HIV on ART (n = 20) | p value |
| --- | --- | --- | --- |
| Demographics, median (IQR) |  |  |  |
| Age (years) | 49 (27-54) | 51 (47-56) | 0.2047 |
| Male, n (%) | 28 (61) | 14 (70) |  |
| Median DNA copies/CD4 T-cell | 175 (168-178) | 173 (165-178) | 0.4356 |
| Weight (kg) | 79 (66-88) | 86 (70-91) | 0.0162 |
| BMI | 25 (23-27) | 27 (23-30) | 0.4345 |
| Laboratory measures, median (IQR) |  |  |  |
| Current CD4 T cell count (cells/uL) | NA | 602 (441-843) | NA |
| >500, n (%) | NA | 13 (65) | NA |
| 200-500, n (%) | NA | 4 (20) | NA |
| <200, n (%) | NA | 3 (15) | NA |
| VL current | NA | 20 (20-48) | NA |
| Medications |  |  |  |
| Years on ART, median (IQR) | NA | 7 (2-14) | NA |

We characterized T cell activation using flow cytometry by measuring CD38 and HLADR co-expression in memory populations of CD8+ T cells (**Figures 1Sa** and **b**). CD38 and HLADR co-expression is a distinguishing feature of intermediate activation and mucosal CD8+ TRM are known to remain in a semi-activated state at baseline^24, 25^ In HIV-uninfected individuals, we observed significantly increased activation of CD8+ T cells in the colon relative to the blood (**Figure 1a** and **b**). In addition, there was also an increase in CD8+ T cell activation in both compartments in ART-treated PWH when compared to HIV-uninfected donors (**Figure 1b**). It has been reported that microbial translocation as measured by soluble CD14 (sCD14) in PWH is linked to systemic inflammation^26^. We observed that the concentration of soluble sCD14 in plasma was increased in HIV infected individuals (**Figure S1c**) correlated with colonic CD8+ T cell activation irrespective of HIV infection status (**Figure 1c**). HIV DNA and RNA from colon showed a median of 273.92 HIV DNA copies/10^6^ CD4+ T cells and a median of 319.09 copies of HIV RNA/10^6^ CD4+ T cells in the colon (**Table 2**). These intracellular HIV DNA and RNA content demonstrate low level HIV replication in ART-treated PWH and potential colon HIV reservoirs. It is plausible that the presence of low-grade HIV replication could drive CD8+ T cell activation in the colon. However, bystander CD8+ T cell activation in ART-treated PWH has been also shown to be HIV antigen-independent^27, 28^. Thus, despite ART treatment, CD8+ T cells in PWH remain aberrantly activated and in particular, higher frequencies of hyperactivated colonic CD8+ T cells are observed when compared to their peripheral counterparts from PWH.

**Figure 1.**
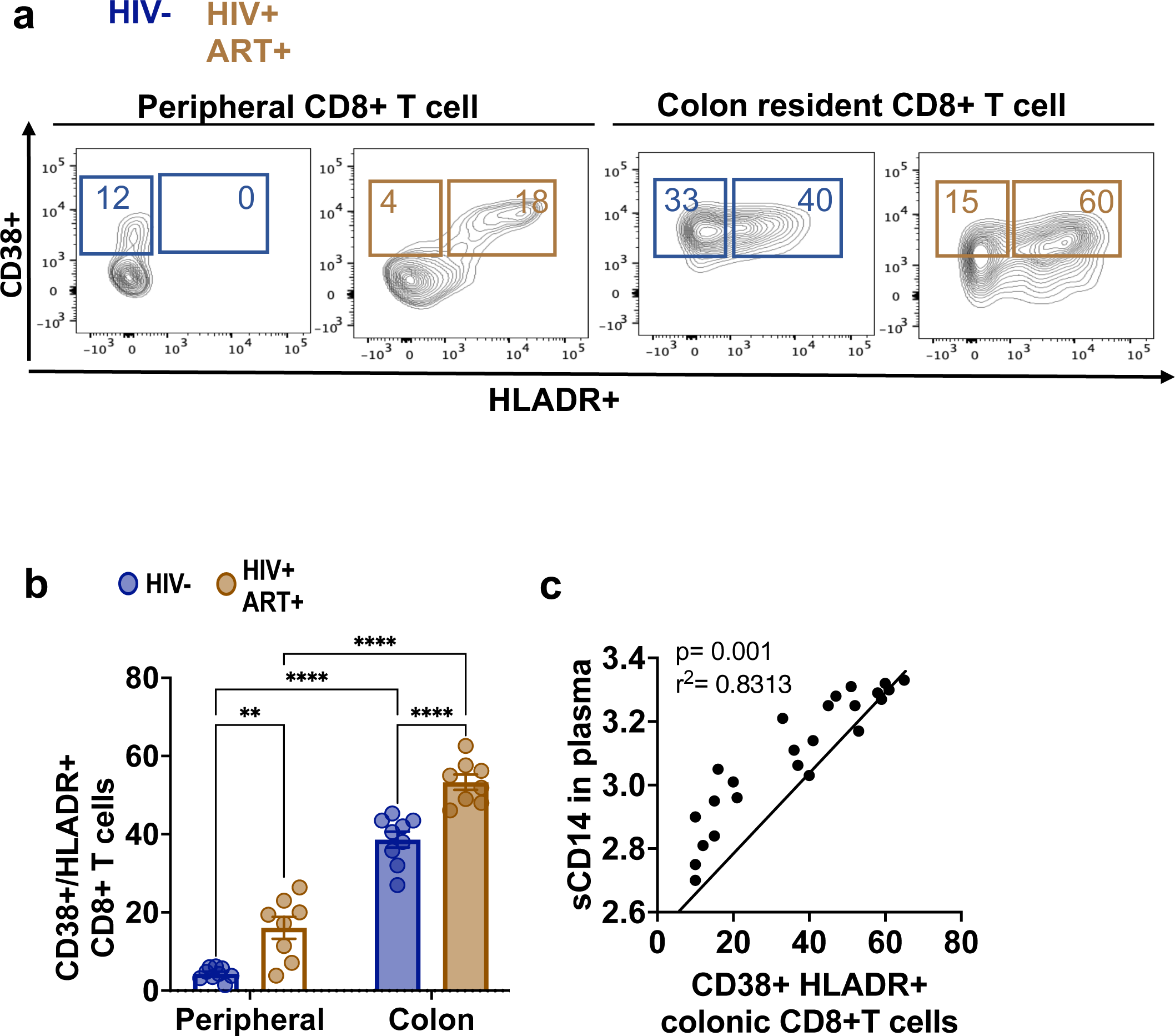
Persistent colon resident CD8+ T cell activation is correlated with systemic inflammation in PWH on ART.

a. Flow cytometric analysis of peripheral and colon resident CD8+ T (CD45RO+) cells showing activation markers CD38+ and HLADR+/CD38+ populations in HIV uninfected healthy donors and PWH on ART.
b. Quantification of peripheral and colon resident CD8+ T (CD45RO+) cells HLADR+/CD38+ populations in HIV uninfected healthy donors and PWH on ART.
c. Correlation between sCD14 and colonic CD8+ T cell activation independent of HIV status.

Table 2.

Table 2
|  | Colon |  |
| --- | --- | --- |
|  | HIV DNA/M CD4 T cells | HIV RNA/M CD4 T cells |
|  | 112.39 | 273.66 |
|  | 65.04 | 6.86 |
|  | 93.74 | 184.29 |
|  | 242.26 | 172.27 |
|  | 120.62 | 3.43 |
|  | 604.71 | 476.15 |
|  | 199.41 | 20.70 |
|  | 437.08 | 1,854.06 |
|  | 156.86 | 63.82 |
|  | 154.76 | 18.87 |
|  | 65.40 | 119.12 |
|  | 78.00 | 100.00 |
|  | 255.23 | 36.67 |
|  | 42.18 | 5.92 |
|  | 169.94 | 7.14 |
|  | 558.18 | 945.43 |
|  | 233.18 | 379.38 |
|  |  | 36.00 |
|  | 865.26 | 440.18 |
|  | 554.27 | 735.58 |
| Median | 263.61 | 109.56 |

### Dysfunction of colon CD8+ TRM from PWH on ART

In peripheral blood, overactivation of CD8+ T cells leads to T cell exhaustion and ultimately alters normal T cell function^29^. T cell dysfunction prevents cells from mounting an appropriate immune response and leads to memory T cell senescence, where memory T cells fail to expand upon restimulation^30–32^. However, this has not been well characterized in the intestine of PWH. We therefore isolated CD8+ T cells from the colon and blood of PWH and HIV-uninfected donors and treated them *ex vivo* with phytohemagglutinin (PHA; **Figure 2a**). We analyzed Ki67, a marker of cellular proliferation^33^, and showed that memory CD8+ T cells (CD8+/CD45RO+) from both the colon and periphery of ART-treated PWH were less proliferative than those memory cell populations from HIV-uninfected donors, although this difference was much greater in the colon than in blood (**Figure 2b**). CD8+ T cells mediate their effector function by secreting cytotoxic cytokines such as IFNψ, TNFα and cytolytic enzymes such as perforin and granzyme B^34^. We showed that when stimulated *ex vivo*, colonic CD8+ TRM from PWH expressed significantly lower intracellular IFNψ than HIV-uninfected donors (**Figure 2c** and **d**). However, peripheral memory CD8+ T cells from PWH have unchanged intracellular IFNψ expression in comparison to those from HIV-uninfected donors, indicating intact effector function capacity. Colonic CD8+ TRM expressed 5-fold lower intracellular IFNψ than their counterparts in periphery (**Figure 2d**), thus demonstrating a suboptimal effector immune response. Similar results were observed for TNFα (**Figure S2a**) and granzyme B secretion (**Figure S2b**). Moreover, HIV-antigen specific CD8+ TRM from the colon were significantly lower than those in the periphery in PWH, as observed in an ELISPOT assay (**Figure S2c**). Studies have shown that HIV infection drives T cell exhaustion of HIV-specific CD8+ T effector cells^30, 31^ and senescence of memory T cells in the periphery of PWH on ART^35, 36^. CD57 which is a marker of senescent CD8+ T cells^37^, showed increased expression on colonic CD8+ TRM from PWH on ART compared to those from HIV-uninfected individuals, with no difference in the periphery (**Figure 2e**). Taken together, these data indicated that colonic CD8+ TRM had overall higher levels of baseline activation than those in the blood and had greater dysfunction in PWH on ART than compared to the periphery and compared to those from HIV-uninfected individuals.

**Figure 2.**
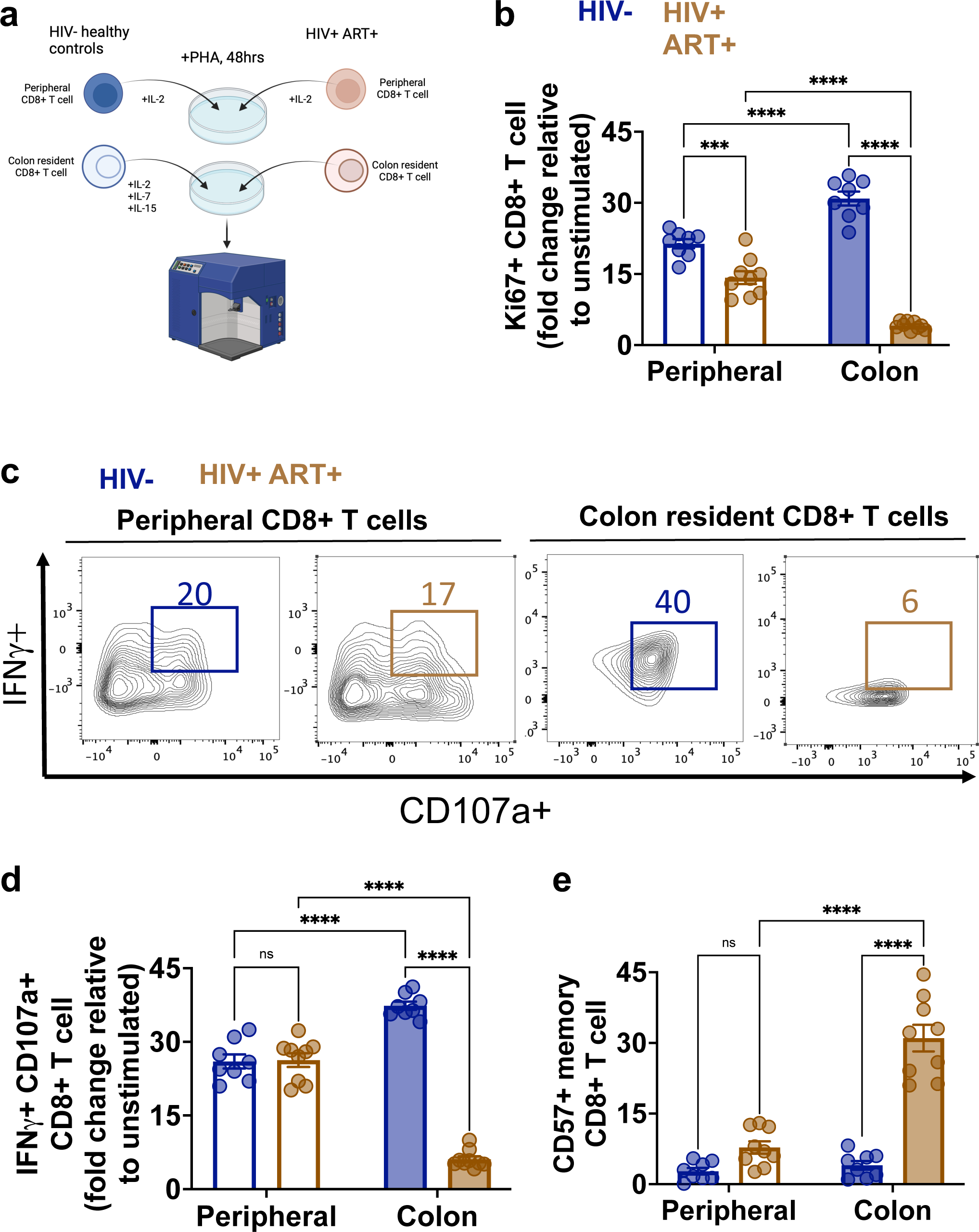
Functional impairment of colon resident CD8+ T cell in PWH on ART.

a. Schematic showing PHA mediated stimulation of peripheral and colon resident CD8+ T (CD45RO+) cell *ex vivo*.
b. Quantification of peripheral and colon resident CD8+ T (CD45RO+) cell in HIV uninfected healthy donors and PWH on ART for proliferative population (KI67+) post stimulation ex vivo.
c. Flow cytometric analysis of peripheral and colon resident CD8+ T (CD45RO+) cells showing intracellular IFNg+ and CD107a+ populations in HIV uninfected healthy donors and PWH on ART.
d. Quantification of peripheral and colon resident CD8+ T (CD45RO+) cell in HIV uninfected healthy donors and PWH on ART for IFNg+/CD107a+ populations.
e. Quantification of senescent marker CD57+ CD8+ T (CD45RO+) cell in peripheral and colon of HIV uninfected healthy donors and PWH on ART

### Mitochondrial function is dysregulated in colon CD8+ TRM in PWH on ART

Previous studies have shown that mitochondrial dysfunction exists in peripheral CD8+ T cells from PWH^19, 21, 23, 38, 39^. Since there are compartmental differences in the mitochondrial metabolism of T cells between the intestines and peripheral blood^40^, understanding the impact of mitochondrial function on colonic CD8+ TRM from PWH is critical. Given the significantly greater degree of CD8+ T cell dysfunction in PWH on ART observed in the colon, we sought to better understand the potential role of mitochondrial function in this defect. We performed flow cytometry of autologous patient colon resident and peripheral CD8+ T cells to measure membrane potential and mitochondrial mass (**Figure S3a**). Mitochondrial membrane potential (MMP; Δψm) refers to the electrical potential difference across the mitochondrial inner membrane, which is generated by the electron transport chain during OXPHOS^13–16^. This potential is critical for the production of ATP, the main energy source of the cell. MitoRed is a cationic dye that accumulates within the mitochondrial matrix in a manner dependent on the magnitude of the MMP. As such, changes in MMP can be detected by changes in MitoRed fluorescence intensity^41^. We found that colonic CD8+ TRM from PWH on ART displayed hyperpolarized mitochondrial membranes relative to HIV-uninfected participants in both the periphery and colon (**Figure 3a**). The MMP of colonic CD8+ TRM is 3-fold higher than autologous peripheral memory CD8+ T cells in ART-treated PWH (**Figure 3a**). This increase in mitochondrial membrane potential of CD8+ T cell is negatively correlated with IFNψ production of CD8+ T cell independent of HIV status (**Figure S3b**), suggestive of mitochondrial MMP induced CD8+ T cell function. Mitochondrial mass has been shown to govern memory T cell activity, where increased mitochondrial mass accompanies T cell activation and differentiation during infection^19, 42, 43^. We measured mitochondrial mass by determining the mean fluorescence intensity (MFI) of MitoGreen, that accumulates in the mitochondrial matrix in an MMP-independent manner^44^. We found no difference in mitochondrial mass in colonic and peripheral CD8+ T cells from PWH when compared to HIV-uninfected donors (**Figure 3b**). However, we showed that *ex vivo* stimulation of colonic CD8+ TRM from HIV-uninfected participants results in increased mitochondrial mass compared (**Figure S3c**). The failure to induce mitochondrial biogenesis in activated CD8+ T cells has also been associated with T cell dysfunction^13, 15, 19^ and these results suggest that CD8+ TRM from PWH on ART fail to increase mitochondrial mass despite being hyperactivated.

**Figure 3.**
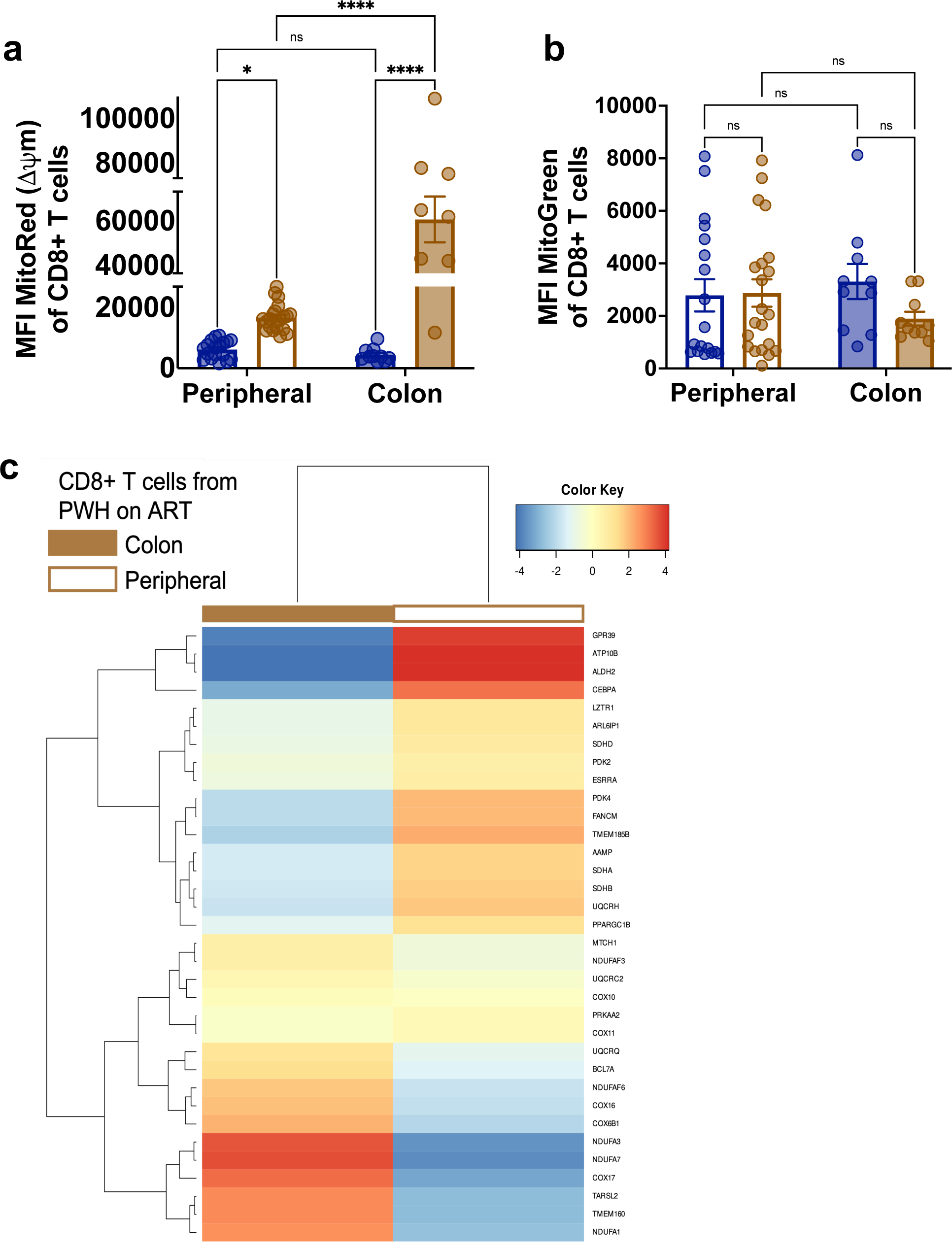
Mitochondrial dysfunction in colon resident CD8+ T cells in PWH on ART.

a. Quantification of MitoRed MFI representing mitochondrial membrane potential (Δψm) of peripheral and colon resident CD8+ T (CD45RO+) cells in HIV uninfected healthy donors and PWH on ART.
b. Quantification of MitoGreen MFI representing mitochondrial mass of peripheral and colon resident CD8+ T (CD45RO+) cells in HIV uninfected healthy donors and PWH on ART.
c. Heatmap representing differential gene expression of mitochondrial OXPHOS related genes between peripheral and colonic CD8+ T cells from PWH on ART.

To further study CD8+ T cell metabolic function, matched CD8+ T cells were isolated from the colon and peripheral blood of ART treated PWH and HIV-uninfected donors for transcriptional analysis. We demonstrated enrichment of genes associated with OXPHOS for both compartments in PWH when compared to HIV-uninfected donors (**Figure S3d and e**). OXPHOS determines mitochondrial ATP production and is dependent on the proper functioning of the electron chain transport (ETC)^13^. The ETC is composed of five complexes embedded in the inner mitochondrial membrane^45^. When we examined CD8+ T cells from the colon and peripheral blood, specific ETC complex-associated genes were decreased in both compartments in PWH on ART compared to HIV-uninfected individuals. We specifically saw genes related to Complex-I of the ETC such as NDUFA family genes (NDUFAF3, NDUFAF6, NDUFA3, NDUFA7. NDUFA1) were downregulated in peripheral CD8+ T cells from PWH, whereas Complex-II of the ETC such as SDH family genes (SDHA, SDHB, SDHD) were downregulated in colonic CD8+ TRM from PWH (**Figure 3c**). This indicated tissue compartment specific alterations in mitochondrial response in PWH. Overall, the hyperpolarized mitochondrial membrane potential in CD8+ T cells and the failure to enhance mitochondrial mass although they remained in activated state in ART-treated PWH were indicative of bioenergetically dysfunctional mitochondria. We also reveal tissue specific compartmental differences in mitochondrial response in colonic and peripheral CD8+ T cells in PWH on ART.

### Impaired mitochondrial metabolism in colon CD8+ TRM in ART-treated PWH

The findings that colonic CD8+ TRM from ART-treated PWH remained in a hyperactivated state, but with defective effector functions, hyperpolarized mitochondrial membrane potential, and impair mitochondrial biogenesis, led us to hypothesize that mitochondrial metabolism would be affected in colonic CD8+ TRM from ART-treated PWH. A metabolically active cell would maintain their mitochondrial basal respiration to generate ATP for their maintenance^46^ and during infection or when activated should increase their mitochondrial respiration to meet the increased demands of ATP^47^. The metabolic profile of peripheral and colon CD8+ T cells was performed using the Mito Stress test with the Seahorse XFe96 Bioanalyzer (**Figure 4a** and **b**) where oxygen consumption was measured, with and without the addition of Oligomycin (to block ATP synthesis), FCCP (to uncouple ATP synthesis from the ETC and trigger maximal oxygen consumption) and Rotenone and Antimycin A (RAA; to inhibit complex I and III of ETC)^48^. Upon FCCP challenge, functional mitochondria from CD8+ T cells accelerate oxygen consumption to mount a strong immune response^48^. Oxygen consumption rate (OCR without Oligomycin, FCCP and RAA is known as the basal respiration) was 3-fold increase in peripheral CD8+ from ART treated PWH than HIV-uninfected donors (**Figure 4c**). Meanwhile, we observed no difference in basal respiration between colonic CD8+ TRM from PWH and HIV-uninfected donors. The failure to increase their mitochondrial respiration though they were in activated state, indicated their metabolic dysfunction. We also measured the respiratory capacity of CD8+ TRM. The respiratory reserve capacity is the difference between maximal respiration and basal respiration. Greater respiratory reserve capacity is indicative of metabolic fitness and considered important for long-term cell T cell survival and function^49, 50^. We found no difference in reserve capacity between peripheral CD8+ T cells from people with HIV and HIV-uninfected donors (**Figure 4d**) suggesting mitochondrial fitness in peripheral CD8+ TRM in PWH on ART. Respiratory reserve capacity was severely impaired in colonic CD8+ TRM from ART treated PWH in compared to HIV-uninfected donor derived TRM (**Figure 4d**) indicating mitochondrial damage. This defect may be indicative of mitochondrial metabolic instability in these cells, which could impact their immune function and potentially contribute to the pathogenesis of HIV.

**Figure 4.**
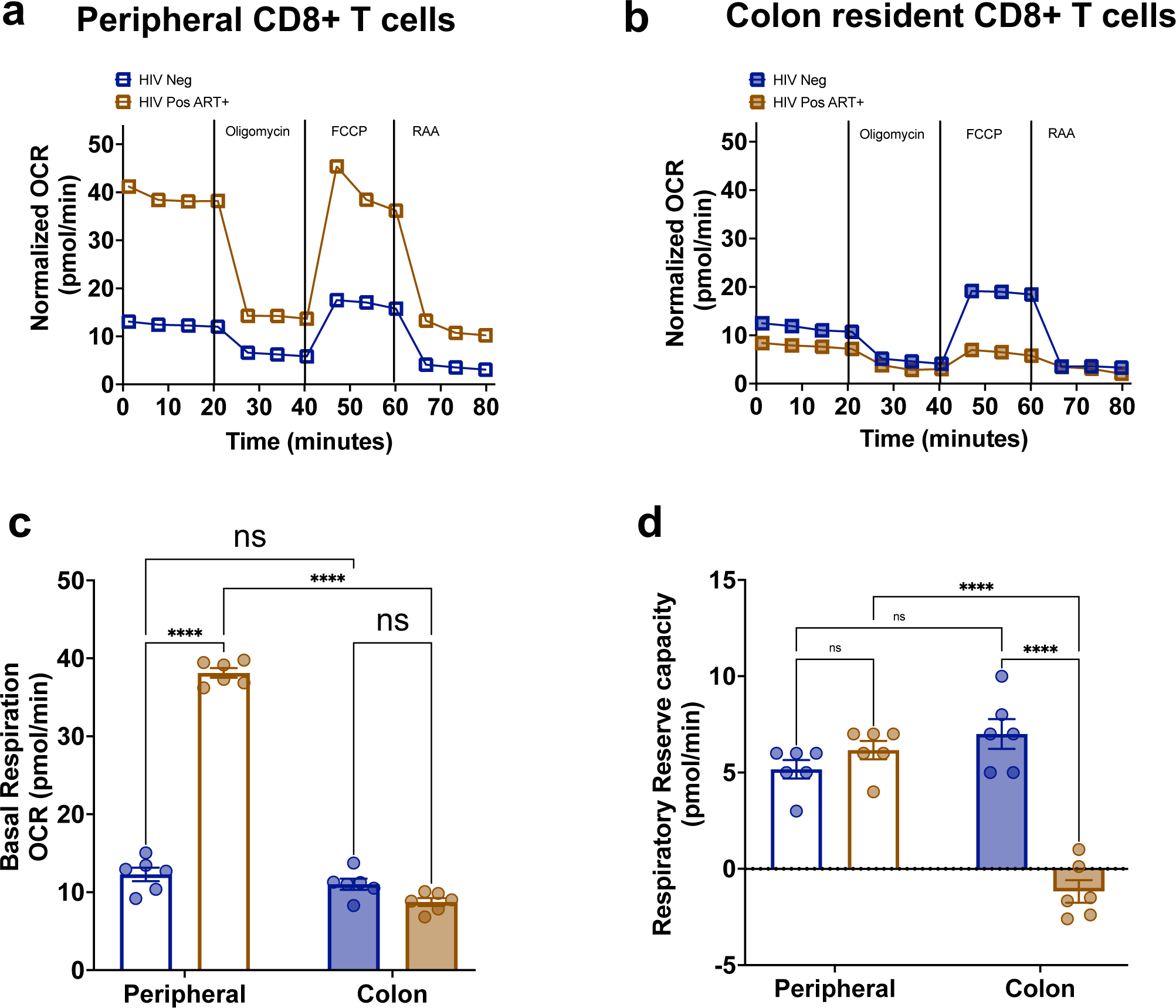
Dysregulation of mitochondrial energy production in colon resident CD8+ T cells in PWH on ART.

a. Real-time oxygen consumption rate (OCR) of peripheral CD8+ T (CD45RO+) cells from HIV uninfected healthy donors and PWH on ART by Seahorse XF analyzer.
b. Real-time oxygen consumption rate (OCR) of colonic CD8+ T (CD45RO+) cells from HIV uninfected healthy donors and PWH on ART by Seahorse XF analyzer.
c. Quantification of Baseline OCR (Basal respiration) of peripheral and colon resident CD8+ T (CD45RO+) cell in HIV uninfected healthy donors and PWH on ART.
d. Quantification of Reserve respiratory capacity (difference of OCR between maximal and basal respiration) of peripheral and colon resident CD8+ T (CD45RO+) cell in HIV uninfected healthy donors and PWH on ART.

ETC Complex II is known to be the primary source of respiratory reserve capacity^52^ and defect in complex II in colonic CD8+ TRM from ART-treated PWH (**Figure S4a**) highlights to the findings in **Figure 4d**. Although there is a decrease in the expression of genes associated with complex-I of the electron transport chain (ETC) in peripheral CD8+ memory T cells from people living with HIV (PWH) (**Figure 3c and S4b**), the impact of this downregulation on the mitochondrial function of these cells is not fully understood. Further experiments are required to elucidate the relationship between complex-I expression and mitochondrial function in CD8+ memory T cells. It is important to conduct additional studies to gain a more comprehensive understanding of the mechanisms underlying this phenomenon and its potential implications for the health and well-being of PWH.

## DISCUSSION

This study characterizes immunological changes associated with GI barrier dysfunction in a cohort of PWH on suppressive ART. Despite the ability of ART treatment to suppress HIV replication in blood, studies have shown that the GI mucosa serve as a reservoir for latent HIV replication^5–8^. Colon CD8+ TRM play a critical role in maintaining intestinal homeostasis between immune activation and tolerance^5, 53^. These TRM have the potential to control HIV replication and reduce the size of the viral reservoir, however, they are not able to eliminate HIV from the tissue^11, 54^. Upon comparative analysis of autologous colon and blood samples from PWH on ART and HIV-uninfected individuals, we reported that the memory subset of colon CD8+ TRM displayed poor memory cell expansion capacity and diminished cytokine secretion relative to their peripheral counterparts and also in PWH on ART relative to those who are uninfected. Additionally, there was significant dysfunction in the mitochondrial bioenergetics and metabolism of colonic CD8+ TRM in PWH on ART.

Most studies in PWH have focused on the findings from peripheral CD8+ T cells, leaving the field with an incomplete understanding of HIV pathophysiology in GI mucosa. We investigated the function of intestinal CD8+ T cells in PWH on ART by leveraging patient intestinal biopsy samples and autologous blood from our cohort of PWH on ART and matched HIV-uninfected control subjects. Our study highlights immune aberrations in the colon of PWH on ART and unique defection in T cell function that are compartmentalized in the intestine. HIV infection promotes cytokine secretions that triggers CD8+ T cell expansion and differentiation^53^. IL-15 released from dendritic cells is a strong promoter of bystander CD8+ T cell activation in PWH independent of active HIV infection, particularly in the mucosal tissues^28^. Colonic CD8+ T cells are continually exposed to microbial stimuli while maintaining a semi-activated state, converting to a fully activated state only upon antigen exposure^25^. The presence of low-grade HIV RNA may be stimulatory to colonic CD8+ TRM even while on ART. The prolonged CD8+ T cell activation may result in immune dysfunction in the form of immunosenescence. Senescence is characterized by cell-cycle arrest that leads to impeded function while remaining metabolically viable^59^. Importantly, this dysfunction does not appear to be limited to HIV-specific memory CD8+ T cells and was also observed in all CD8+ TRM residing in the colon of PWH on ART. Although advances over the past decade have shed considerable light on the mechanisms of T cell senescence and have revealed novel therapeutic opportunities to reverse T cell dysfunctional states, we still understand surprisingly little about the molecular mechanisms by which these pathways operate.

Mitochondrial dysfunction has been linked with cellular senescence^60, 61^; however, it is unclear whether mitochondria play a causal role. Our studies show that colonic CD8+ T cells from PWH on ART are dysregulated in comparison to their counterparts in the periphery. Although CD8+ T cell activation is linked to mitochondrial biogenesis during infection^43^, the mitochondrial mass of persistently activated colonic CD8+ TRM remains unchanged in PWH on ART. While mitochondrial depolarization (decrease in Δψm) is observed during homeostasis, hyperpolarization (increase in Δψm) is often associated with CD8+ T cell dysfunction and chronic inflammation^60–62^. Hyperpolarized colonic CD8+ T cells are indicative of immune aberration in PWH on ART. While mitochondrial metabolic activity plays a central role in regulating T cell activation and proliferation, dampened mitochondrial metabolism, as shown by the diminished mitochondrial respiration of colonic CD8+ T cells in PWH on ART, emphasizes their immunosenescence together with CD57 expression. The enhanced mitochondrial function of peripheral CD8+ T cells from PWH on ART relative to the colonic CD8+ T cells illustrates their improved functioning upon ART treatment. However, the transcriptomic downregulation of Complex-I of the ETC in peripheral CD8+ T cells require further investigation to understand its specific role. Among mitochondrial respiratory parameters, the SRC represents a particularly robust functional parameter to evaluate mitochondrial reserve^49^. Mitochondrial SRC of a cell is a critical component of its bioenergetics that can be utilized during an increase in energy demand. During metabolic adaptation, a cell draws on mitochondrial reserve to meet increasing needs^49, 50^. Downregulation of SRC in colonic CD8+ T cells from PWH on ART is indicative of their dampened SRC. This may contribute to impaired colonic CD8+ T cells immune responses in PWH on ART. Since Complex-II of ETC is a contributor of the SRC^52^, the reduction of complex II of the ETC in colonic CD8+ T cells refers to a decrease in the activity of the enzyme complex II, which is part of the ETC responsible for generating ATP (adenosine triphosphate) through oxidative phosphorylation. This reduction suggests that the CD8+ T cells in the colon are experiencing some form of stress or impairment, which could be due to the effects of HIV infection or antiretroviral therapy. On the other hand, the fairly increased complex I activity refers to an increase in the activity of complex I of the ETC. This complex is responsible for initiating the electron transport process in the ETC. This increase in activity could be an adaptive mechanism to compensate for the reduction in complex II activity. Overall, these changes suggest that the CD8+ T cells in the colon of PWH on ART are undergoing adaptive changes to maintain their energy production and survive in the hostile environment of the GI tract. However, these changes may also affect the cells’ ability to effectively control the HIV infection, which is why boosting mitochondrial activity is a potential therapeutic strategy to enhance the immune response against the virus.

To our knowledge, this is the first report examining the characteristics and functionality of patient colon resident and peripheral CD8+ T cells in ART-treated PWH. The hyperactivation of colonic CD8+ TRM play an important role in disrupting GI mucosal homeostasis in PWH on ART. While mitochondrial dysregulation is a strong predictor of immune dysfunction in PWH, the mechanism remains elusive. Overall, these findings highlight that tissue specific studies on the impact of HIV infection are a necessary complement to studies of circulating immune cells. Future mechanistic and clinical studies are required to develop the next generation of tissue targeted immune-based interventions in PWH on ART to improve overall immune function.

## METHODS

### Study participants and sample collection

The study contained two populations: 20 PWH on ART treatment with undetectable viremia and 46 age-matched uninfected, HIV-uninfected control subjects. Participant characteristics are summarized in the **Table 1**. Sigmoid colon pinch biopsies with participant-matched blood samples were obtained during upper and lower endoscopy. Blood samples were obtained from HIV uninfected, HIV-uninfected controls. Clinical information, including CD4+T cell count and median years on ART, were collected through the Ragon Institute Cellular Immunology Database Platform. In **Table 2**, we compare patient autologous colon and peripheral blood HIV DNA and RNA in CD4+T cells from PWH.

### Markers of microbial translocation and inflammation

The concentration of some of the plasma markers was measured by ELISA using Human sCD14 DuoSet ELISA (R&D Systems, DY383) and Human FABP2/I-FABP DuoSet ELISA (R&D Systems, DY3078), per manufacturer’s instructions.

### Blood and Tissue processing

#### PBMC isolation

Peripheral blood mononuclear cells (PBMCs) were isolated from the blood of HIV uninfected and HIV infected ART-treated patients by density gradient centrifugation. Blood was transferred to a 50 mL conical and centrifuged at 2600 rpm for 15 minutes. The resultant plasma layer was discarded, and the remainder was volumed up to 30mL with Hank’s Balanced Salt Solution (HBSS). This mixture was then layered with 15mL of Histopaque-1077, after which the conical was spun at 1500 rpm for 45 minutes. The PBMC layer was then collected in a new tube, washed twice with HBSS, counted, and frozen in a 10% DMSO solution.

#### Colon mucosal lymphocyte isolation

Lymphocytes were isolated from intestinal biopsy pinches by modifying the protocol as described^63–65^.

#### IEL isolation

Biopsy pinches were transferred to a 50mL conical containing IEL stripping buffer (PBS, 10mM DTT, 5mM EDTA, 10mM HEPES, & 5% FCS) and incubated under continuous rotation for 20 minutes at 37°C. After separating tissue pieces from the supernatant, the tissue was again treated with IEL buffer and incubated while the supernatant was spun down and IELs were collected. This process was repeated until all IELs had been isolated from the digested biopsy tissue.

#### LP isolation

Following IEL isolation, biopsy pinches were placed into a Collagenase digestion solution. Tissue pinches were incubated in Collagenase at 37°C for one hour to release lamina propria cells. These cells, in combination with isolated IELs, were enriched for CD45+ cells by positive immunomagnetic selection using Miltenyi CD45 microbeads.

### Flow cytometry

Cryopreserved PBMCs and colon mucosal lymphocytes were thawed, washed, and counted. 100,000 cells per sample were allotted into FACS tubes, along with 20,000 cells for each fluorescence minus one (FMO) control and mitochondrial dye compensation control tube. FMOs were created for each surface antibody and mitochondrial dye. The following fluorescence-tagged antibodies were used for PBMCs: CD3-BUV737/PerCP-Cy5.5, CD8-BUV395/V500, CD45 RO-APC and LIVE/DEAD Fixable Blue Stain. GI mucosal lymphocytes were stained with the fluorescence-tagged antibodies: CD3-BUV737/PerCP-Cy5.5, CD8-BUV395/V500, CD103-PECy7, S1PR1-PE, CD45 RO-APC and LIVE/DEAD Fixable Blue Stain. All cells were incubated with Fc block for 5 minutes at room temperature (RT) and stained with surface antibodies for 20 minutes at RT. For intracellular staining, all cells were incubated with MitoTracker Red (2nM) and MitoTracker Green (2nM) for 45 minutes at 37°C. After washing, cells were processed by the flow cytometer (4 Laser LSR II). Compensation controls for each surface antibody were prepared using anti-mouse IgG compensation beads. Separate application settings were created in BD FACSDiva for PBMCs and colon mucosal lymphocytes and used in each experiment for consistent fluorophore excitation. Data were analyzed with FlowJo 10.8.1 using a hierarchical gating strategy. Gating strategies for PBMCs and colon mucosal lymphocytes are elaborated in Supplementary Figure 1c and d, respectively.

### Stimulation of lymphocytes and Intracellular Cytokine Staining

#### PHA stimulation

Colon mucosal lymphocytes from 10 uninfected subjects were stimulated with 4ug/mL of Phytohemagglutinin (PHA) in 24-well tissue culture plates at 37°C. One million cells were plated into each well, with a total of two technical replicates for each stimulated sample and unstimulated control. Cells were cultured in complete RPMI media (10% FBS) in the presence of 50 IU/mL human IL-2, 10 ng/mL human IL-7, and 10 ng/mL human IL-15. After 48 hours, cells were harvested and processed by flow cytometry.

#### Antigen stimulation

For the antigenic stimulation assay, cells were incubated at 1×106 cells per 200μL complete RPMI media for 6h in the presence of, anti-CD107a, 1μM GolgiStopTM (BD Biosciences), brefeldin A (5μg/mL, Sigma Aldrich) and Gag-peptide pool (3.5μg/ mL), or media containing DMSO (peptide solvent) as negative control. Cells were incubated for 10 minutes in PBS with 0.5mM EDTA prior to staining for surface markers and viability (Blue Viability dye, Invitrogen) for 20 minutes at room temperature. Cells were fixed in 4% paraformaldehyde (PFA) and permeabilized using FACS Perm 2 (BD Biosciences) prior to intracellular staining for CD3-PerCP-Cy5.5, CD45RO-APC, CD8-V500 and IFN-γ-Pe-Cy7. The Transcription Factor Buffer Set (BD Biosciences) was substituted for 4% PFA and FACS Perm 2 when staining for transcription factors. In such cases, cells were fixed with Transcription Factor Fix/Perm Buffer for 40 minutes at 4°C, washed twice in Transcription Factor Perm/Wash Buffer, and stained with antibodies for 40 minutes at 4°C in Transcription Factor Perm/Wash. For all protocols, after staining, cells were re-suspended in 1% PFA and stored at 4°C in the dark until analysis within 24h.

#### IFN-gamma ELISpot assay

IFN-γ ELISpot assays were performed using the Human IFN-γ ELISpot PLUS ALP kit (Mabtech) according to the manufacturer’s protocol with some modifications. Basic assay pipetting workflow was performed as follows; the assay plate was re-hydrated by washing with 1X PBS and blocked with CTL-Test media (CTLT-005, Cellular Technology Limited) for 2–6 hours at RT. Meanwhile, the stimuli (gag-peptide pool) were prepared at 2X and, to each well of a blocked ELISpot plate, 100 µl of 2X stimuli was added in duplicate for each PBMC and colon lymphocyte sample to be tested. Briefly, the 2X concentrations of all stimuli were prepared as follows: peptide pool stimuli were diluted to either 4 µg/mL for the PHA positive control, 0.2–0.5% DMSO for the unstimulated negative control, and CTL-Test media alone for the background negative control (containing medium only without any cells). Freshly thawed PBMCs and colon lymphocytes were thawed quickly at 37 °C and washed in CTL-Wash media (CTLW-010, Cellular Technology Limited) supplemented with 50 U/mL Pierce Universal Nuclease (88701, ThermoFisher). Lymphocytes, with a viability ≥70%, were diluted in CTL-Test media to a concentration of 1.0 × 10^6^ cells/mL and plated 100 µl/well in duplicate for each stimulus (1.0 × 10^5^ cells/well). Lymphocytes were stimulated for 18–24 h at 37 °C + 5% CO2 after which cells were removed and plates were incubated with a streptavidin alkaline phosphatase-coupled detection antibody for 1 h. Spots were visualized after incubation with the substrate BCIP/NBT-plus for 3 min ± 15 s at RT. Plate development was stopped with a water wash and the plate was air-dried overnight at RT, avoiding exposure to light. Spots were enumerated using an automated spot counter (ImmunoSpot® CTL S6 Micro Analyzer; Cellular Technology Limited) within 24–96 h and spot counts are further analyzed as spot forming units (SFU) per well. The Immunospot settings used were a sensitivity of 145 or 187, background balance of 10, spot separation of 1–3, and spot size minimum of 0.0015 to maximum of 9.6296 or 9.6466 mm^2^. These parameters were optimized for resolution and accurate enumeration of spots. Raw data included images for each well for visual quality checking and the corresponding spot forming unit (SFU) counts per well.

### Measurement of blood and colon HIV-1 DNA and RNA using ddPCR

Total PBMC and colon tissue mononuclear cells from study participant were subjected to DNA extraction using DNeasy Blood & Tissue Kits (QIAGEN). Total HIV-1 DNA and host cell concentrations in the DNA extracts were estimated using Bio-Rad ddPCR, using primers and probes covering HIV-1 5′ LTR-gag HXB2 coordinates 684–810 (forward primer 5′-TCTCGACGCAGGACTCG-3′, reverse primer 5′-TACTGACGCTCTCGCACC-3′ probe/56-FAM/CTCTCTCCT/ZEN/TCTAGCCTC/31ABkFQ/, and human RPP30 gene (forward primer 5′-GATTTGGACCTGCGAGCG-3′, reverse primer 5′-GCGGCTGTCTCCACAAGT-3′, probe/56-FAM/CTGACCTGA/ZEN/AGGCTCT/31ABkFQ/). ddPCR was performed using the following thermocycler program: 95°C for 10 minutes, 45 cycles of 94°C for 30 seconds, and 60°C for 1 minutes, 72°C for 1 minute. The droplets were subsequently read by the Bio-Rad QX100 droplet reader, and data were analyzed using QuantaSoft software (Bio-Rad)^66^.

RNA was extracted using Phenol-Chloroform assay. The following protocol was performed. Brilliant II SYBR Green qPCR kit was used. Reference dye (ROX) was prepared by diluting 1:10,000 with DEPC H2O.

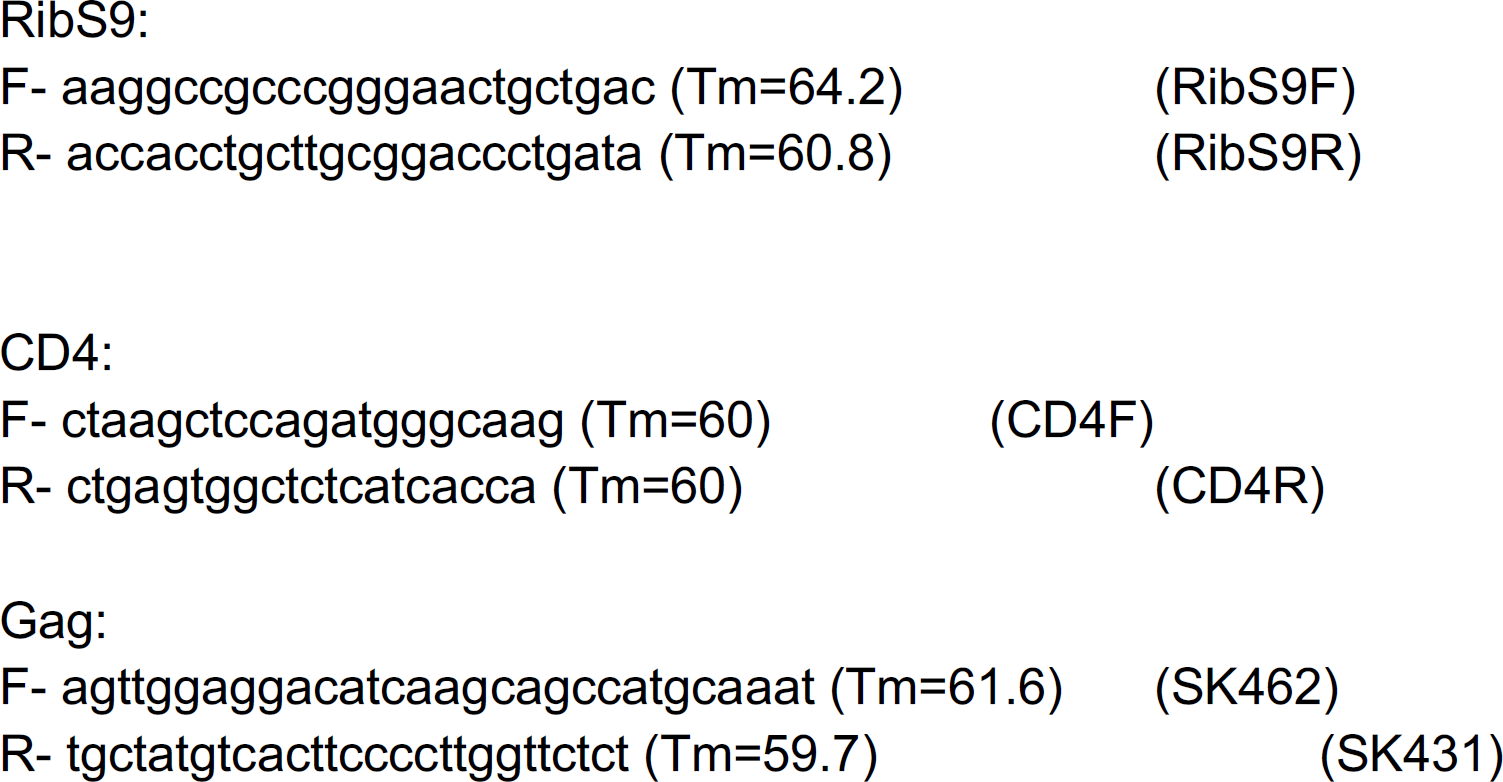

The PCR program used as follows:

95C 10min 95C 00:30

60C 01:00

75C 00:30

(40 cycles)

95C 1:00

55C 00:30

95C 00:30

### Bulk sequencing of CD8+ T cells

#### RNA isolation, library construction, sequencing, and alignment

CD8+ T cells from PBMCs and colon were FACS sorted directly into 50 μL of RLT Lysis Buffer (QIAGEN) supplemented with 1% v/v 2-mercaptoethanol. Briefly, 50 μL of mixed lysate from each sample was transferred to a skirted 96 well plate. Genetic material was pulled down and purified by mixing the lysate in each well with 2.2x volumes of Agencourt RNAClean XP SPRI beads (Beckman Coulter) and washing 3x with 75 μL of 80% ethanol. After drying, the SPRI beads were re-suspended in 4 μL of pre-reverse transcription (RT) mix, incubated for 3 min at 72°C, and placed on ice. Next, Smart-Seq2 Whole Transcriptome Amplification (WTA) was performed: 7 μL of RT mix was added to each well and RT was carried out; then, 14 μL of PCR mix was added to each well and PCR was performed. Thereafter a cDNA cleanup was performed using 0.6x and 0.8x volumes of Agencourt AMPure XP SPRI beads (Beckman Coulter) which was then quantified using a Qubit dsDNA HS Assay Kit (Life Technologies). Library size and quality were measured by Bioanalyzer using a High Sensitivity DNA Analysis Kit (Agilent Technologies). Sequencing libraries were prepared from WTA product using Nextera XT (Illumina). After library construction, a final AMPure XP SPRI clean-up (0.8 volumes) was conducted. Library concentration and size were measured with the KAPA Library Quantification kit (KAPA Biosystems) and a TapeStation (Agilent Technologies), respectively. Finally, samples were sequenced on a NextSeq500 (30 bp paired end reads) to an average depth of 5 million reads. Reads were aligned to hg38 (Gencode v21) using RSEM and TopHat^73^ and estimated counts and transcripts per million (TPM) matrices generated. Any samples with fewer than 5×105 or more than 6×106 aligned reads or fewer than 10,000 uniquely expressed genes were removed from subsequent analysis. *RNA-Seq Differential Expression Analysis:* Differential expression analysis was performed using DESeq2 (v1.18.1)^67^. Expected counts from biological replicates for each cell type and participant were averaged prior to differential expression to prevent participant specific genes from generating false positives and reduce spurious heterogeneity from small (100-cell) populations. Small populations may show skewed expression based on the cell composition within; thus, this replicate averaging approach is particularly important given our limited access to pediatric tissue sources and low frequency of these immune populations in order to remove further bias from small population sorts.

#### Gene Set Analysis

Gene set analysis was performed using Ingenuity Pathway Analysis (IPA; Winter 2019 Release, QIAGEN Inc.) and Gene Set Enrichment Analysis (GSEA) using the piano package in R (1.18.1). For IPA, DEGs whose FDR corrected q < 0.1 were used in the “Core” analysis with the log2FC and q values included in the analysis. To implement GSEA on our DESeq2 results, we used the log2FC of all genes whose FDR corrected q < 0.1 as t-value input into the runGSA function with setting the argument geneSetStat = “gsea.” We chose to use the KEGG and GO databases (downloaded from MSigDB v7.0)^68^ for GSEA analysis as these databases are well annotated for metabolic and cellular activation gene sets that are cell-type agnostic.

### Seahorse mitochondrial metabolic analysis

#### Cell preparation and plating

The Agilent Seahorse XFp Cell Mito Stress Test was performed according to manufacturer’s protocol using the Seahorse XFp Analyzer. Peripheral and GI mucosal CD8+ T cells were purified from PBMCs and GI mucosal lymphocytes, respectively, by positive magnetic cell separation. CD8+ T cells were washed in XF medium, counted, and resuspended in XF medium at a concentration of 5×10^6^ cells/mL. 2×10^5^ cells/well were plated in a 96-well tissue culture plate, centrifuged at 1500rpm for 5 minutes, and covered with 140 μL of XF medium. The plate was placed in a non-CO2 incubator overnight at 37 degrees Celsius.

#### Cartridge Preparation

200μL of calibrant was added to each well of the Seahorse XF Cell Culture Miniplate. The miniplate was coated with Poly-D-lysine and incubated overnight at 37°C in non-CO2 conditions. Cells were transferred to the Miniplate the following day.

#### Program Preparation

On day 2, the Mito Stress test was programmed in the Seahorse console:

1. Calibrate
2. Equilibrate
3. Base Line Readings (3x Loop)
4. Mix 1 → 3 min Wait → 2 min Measure → 3 min End
5. Inject Port B (3x Loop)
6. Mix 2 → 3 min Wait → 2 min Measure → 3 min End
7. Inject Port C (3x Loop)
8. Mix 3 → 3 min Wait → 2 min Measure → 3 min End
9. End

Reagents were prepared at the following concentrations: 1μM oligomycin, 1.5 μM FCCP, 100nM rotenone, and 1μM antimycin A. Drugs were then loaded into the delivery ports of the sensor cartridge with a multichannel pipette:

Port B: 20μL oligomycin

Port C: 22μL FCCP

Port D: 24μL rotenone + antimycin A

## Statistical Analysis

Statistical analysis was performed using GraphPad Prism 9. Each figure legend indicates the statistical test performed. P-values less than 0.05 were taken as significant and represented by asterisk as follows: \**P* ≤ 0.05, \*\**P* ≤ 0.01, \*\*\**P* ≤ 0.001, \*\*\*\**P <* 0.0001.

## Notes

### Competing Interest Statement

The authors have declared no competing interest.

## References

1. McBrien JB, Kumar NA, Silvestri G. Mechanisms of CD8+ T cell-mediated suppression of HIV/SIV replication. Eur J Immunol. 2018 Jun;48(6):898–914. doi: 10.1002/eji.201747172. Epub 2018 Mar 26. PMID: 29427516; PMCID: PMC6531861.

2. FitzPatrick, Michael E B et al. “Human intestinal tissue-resident memory T cells comprise transcriptionally and functionally distinct subsets.” Cell reports vol. 34,3 (2021): 108661. doi: 10.1016/j.celrep.2020.108661

3. Kumar, Brahma V et al. “Human Tissue-Resident Memory T Cells Are Defined by Core Transcriptional and Functional Signatures in Lymphoid and Mucosal Sites.” Cell reports vol. 20,12 (2017): 2921–2934. doi: 10.1016/j.celrep.2017.08.078

4. Brenchley, Jason M et al. “Microbial translocation is a cause of systemic immune activation in chronic HIV infection.” Nature medicine vol. 12,12 (2006): 1365–71. doi:10.1038/nm1511

5. Mehandru, S. et al. Lack of mucosal immune reconstitution during prolonged treatment of acute and early HIV-1 infection. PLoS Med. 2006 3, e484, doi: 10.1371/journal.pmed.0030484.

6. Chun, Tae-Wook et al. “Persistence of HIV in gut-associated lymphoid tissue despite long-term antiretroviral therapy.” The Journal of infectious diseases vol. 197,5 (2008): 714–20. doi:10.1086/527324

7. Belmonte L., Olmos M., Fanin A., Parodi C., Bare P., Concetti H., et al. (2007). The intestinal mucosa as a reservoir of HIV-1 infection after successful HAART. AIDS 21 2106–2108. 10.1097/qad.0b013e3282efb74b

8. Jacobs, Jana L et al. “Persistent HIV-1 Viremia on Antiretroviral Therapy: Measurement and Mechanisms.” Frontiers in microbiology vol. 10 2383. 15 Oct. 2019, doi:10.3389/fmicb.2019.02383

9. Lange, Joshua et al. “Human mucosal tissue-resident memory T cells in health and disease.” Mucosal immunology vol. 15,3 (2022): 389–397. doi:10.1038/s41385-021-00467-7

10. Lyu, Yangbao et al. “An Overview of Tissue-Resident Memory T Cells in the Intestine: From Physiological Functions to Pathological Mechanisms.” Frontiers in immunology vol. 13 912393. 31 May. 2022, doi:10.3389/fimmu.2022.912393

11. van Wijk, Femke, and Hilde Cheroutre. “Mucosal T cells in gut homeostasis and inflammation.” Expert review of clinical immunology vol. 6,4 (2010): 559–66. doi:10.1586/eci.10.34

12. Sáez-Cirión A, Sereti I. Immunometabolism and HIV-1 pathogenesis: food for thought. Nat Rev Immunol. 2021 Jan;21(1):5–19. doi: 10.1038/s41577-020-0381-7. Epub 2020 Aug 6. PMID: 32764670.

13. Hill, Bradford G et al. “Integration of cellular bioenergetics with mitochondrial quality control and autophagy.” Biological chemistry vol. 393,12 (2012): 1485–1512. doi:10.1515/hsz-2012-0198

14. Mills, Evanna L et al. “Mitochondria are the powerhouses of immunity.” Nature immunology vol. 18,5 (2017): 488–498. doi:10.1038/ni.3704

15. Buck MD, O’Sullivan D, Klein Geltink RI, et al. Mitochondrial Dynamics Controls T Cell Fate through Metabolic Programming. Cell. 2016;166(1):63–76. doi: 10.1016/j.cell.2016.05.035

16. Quintana A, Hoth M. Mitochondrial dynamics and their impact on T cell function. Cell Calcium. 2012;52(1):57–63. doi: 10.1016/j.ceca.2012.02.005

17. Huckestein, Brydie R, and John F Alcorn. “Improving Mitochondrial Function in Viral Infection: Targeting Cellular Metabolism.” American journal of respiratory cell and molecular biology vol. 66,6 (2022): 598–600.

18. Chiu, Chris Y et al. “Soluble immune checkpoints as correlates for HIV persistence and T cell function in people with HIV on antiretroviral therapy.” Frontiers in immunologyvol. 14 1123342. 28 Mar. 2023, doi:10.3389/fimmu.2023.1123342

19. Petrovas, Constantinos et al. “Increased mitochondrial mass characterizes the survival defect of HIV-specific CD8(+) T cells.” Blood vol. 109,6 (2007): 2505–13. doi:10.1182/blood-2006-05-021626

20. Deguit, Christian Deo T, et al. “Some Aspects of CD8+ T-Cell Exhaustion Are Associated with Altered T-Cell Mitochondrial Features and ROS Content in HIV Infection.” Journal of acquired immune deficiency syndromes (1999) vol. 82,2 (2019): 211-219. doi:10.1097/QAI.0000000000002121

21. Rahman, Akm Nur-Ur et al. “Elevated glycolysis imparts functional ability to CD8^+^ T cells in HIV infection.” Life science alliance vol. 4,11 e202101081. 21 Sep. 2021, doi:10.26508/lsa.202101081

22. Perdomo-Celis, Federico et al. “Reprogramming dysfunctional CD8+ T cells to promote properties associated with natural HIV control.” The Journal of clinical investigation vol. 132,11 (2022): e157549. doi:10.1172/JCI157549

23. Alrubayyi, Aljawharah et al. “Functional Restoration of Exhausted CD8 T Cells in Chronic HIV-1 Infection by Targeting Mitochondrial Dysfunction.” Frontiers in immunology vol. 13 908697. 5 Jul. 2022, doi:10.3389/fimmu.2022.908697

24. Kestens, L et al. “Expression of activation antigens, HLA-DR and CD38, on CD8 lymphocytes during HIV-1 infection.” AIDS (London, England) vol. 6,8 (1992): 793–7. doi:10.1097/00002030-199208000-00004

25. Konjar, Špela, and Marc Veldhoen. “Dynamic Metabolic State of Tissue Resident CD8 T Cells. Frontiers in Immunology. vol. 10 1683. 17 Jul. 2019, doi:10.3389/fimmu.2019.01683

26. Younas, Mehwish et al. “Microbial Translocation Is Linked to a Specific Immune Activation Profile in HIV-1-Infected Adults with Suppressed Viremia.” Frontiers in immunology vol. 10 2185. 13 Sep. 2019, doi:10.3389/fimmu.2019.02185

27. Perdomo-Celis, Federico et al. “CD8+ T-Cell Response to HIV Infection in the Era of Antiretroviral Therapy.” Frontiers in immunology vol. 10 1896. 9 Aug. 2019, doi:10.3389/fimmu.2019.01896

28. Younes, Souheil-Antoine et al. “IL-15 promotes activation and expansion of CD8+ T cells in HIV-1 infection.” The Journal of clinical investigation vol. 126,7 (2016): 2745–56. doi:10.1172/JCI85996

29. Wherry, E John, and Makoto Kurachi. “Molecular and cellular insights into T cell exhaustion.” Nature reviews. Immunology vol. 15,8 (2015): 486–99. doi:10.1038/nri3862

30. Fenwick, Craig et al. “T-cell exhaustion in HIV infection.” Immunological reviews vol. 292,1 (2019): 149–163. doi:10.1111/imr.12823

31. Day, Cheryl L et al. “PD-1 expression on HIV-specific T cells is associated with T-cell exhaustion and disease progression.” Nature vol. 443,7109 (2006): 350–4. doi:10.1038/nature05115

32. Cobos Jiménez V, Wit FW, Joerink M, Maurer I, Harskamp AM, Schouten J, et al. T-cell activation independently associates with immune senescence in HIV-infected recipients of long-term antiretroviral treatment. J Infect Dis. (2016) 214:216–25. doi: 10.1093/infdis/jiw146

33. Sachsenberg, N et al. “Turnover of CD4+ and CD8+ T lymphocytes in HIV-1 infection as measured by Ki-67 antigen.” The Journal of experimental medicine vol. 187,8 (1998): 1295–303. doi:10.1084/jem.187.8.1295

34. Kiniry, Brenna E et al. “Detection of HIV-1-specific gastrointestinal tissue resident CD8+ T-cells in chronic infection.” Mucosal immunology vol. 11,3 (2018): 909–920. doi:10.1038/mi.2017.96

35. Gangcuangco, Louie Mar A, et al. “Correction: Mitochondrial oxidative phosphorylation in peripheral blood mononuclear cells is decreased in chronic HIV and correlates with immune dysregulation.” PloS one vol. 16,3 e0249428. 25 Mar. 2021, doi: 10.1371/journal.pone.0249428

36. Schank, Madison et al. “The Impact of HIV- and ART-Induced Mitochondrial Dysfunction in Cellular Senescence and Aging.” Cells vol. 10,1 174. 16 Jan. 2021, doi:10.3390/cells10010174

37. Ahmed, Raya et al. “CD57^+^ Memory T Cells Proliferate In Vivo.” Cell reports vol. 33,11 (2020): 108501. doi: 10.1016/j.celrep.2020.108501

38. Butterfield, Tiffany R et al. “Dysfunctional Immunometabolism in HIV Infection: Contributing Factors and Implications for Age-Related Comorbid Diseases.” Current HIV/AIDS reports vol. 17,2 (2020): 125–137. doi:10.1007/s11904-020-00484-4

39. Jimnez V.C., Wit F.W.N.M., Joerink M., Maurer I., Harskamp A.M., Schouten J., Prins M., Van Leeuwen E.M.M., Booiman T., Deeks S.G., et al. T-Cell Activation Independently Associates with Immune Senescence in HIV-Infected Recipients of Long-term Antiretroviral Treatment. J. Infect. Dis. 2016; 214:216–225. doi: 10.1093/infdis/jiw146.

40. Varanasi, Siva Karthik et al. “Determinants of Tissue-Specific Metabolic Adaptation of T Cells.” Cell metabolism vol. 32,6 (2020): 908–919. doi: 10.1016/j.cmet.2020.10.013

41. Perry, Seth W et al. “Mitochondrial membrane potential probes and the proton gradient: a practical usage guide.” BioTechniques vol. 50,2 (2011): 98–115. doi:10.2144/000113610

42. Scharping, Nicole E et al. “The Tumor Microenvironment Represses T Cell Mitochondrial Biogenesis to Drive Intratumoral T Cell Metabolic Insufficiency and Dysfunction.” Immunity. vol. 45,2 (2016): 374–88. doi: 10.1016/j.immuni.2016.07.009

43. Sena, Laura A et al. “Mitochondria are required for antigen-specific T cell activation through reactive oxygen species signaling.” Immunity. vol. 38,2 (2013): 225–36. doi: 10.1016/j.immuni.2012.10.020

44. Monteiro, Lauar de Brito, et al. “Using flow cytometry for mitochondrial assays.” MethodsX vol. 7 100938. 28 May. 2020, doi: 10.1016/j.mex.2020.100938

45. Nolfi-Donegan, Deirdre et al. “Mitochondrial electron transport chain: Oxidative phosphorylation, oxidant production, and methods of measurement.” Redox biology vol. 37 (2020): 101674. doi: 10.1016/j.redox.2020.101674

46. Weinberg, Samuel E et al. “Mitochondria in the regulation of innate and adaptive immunity.” Immunity vol. 42,3 (2015): 406–17. doi: 10.1016/j.immuni.2015.02.002

47. Angajala, Anusha et al. “Diverse Roles of Mitochondria in Immune Responses: Novel Insights Into Immuno-Metabolism.” Frontiers in immunologyvol. 9 1605. 12 Jul. 2018, doi:10.3389/fimmu.2018.01605

48. Ron-Harel, Noga et al. “Mitochondrial metabolism in T cell activation and senescence: a mini-review.” Gerontology vol. 61,2 (2015): 131–8. doi:10.1159/000362502

49. Marchetti, Philippe et al. “Mitochondrial spare respiratory capacity: Mechanisms, regulation, and significance in non-transformed and cancer cells.” FASEB journal: official publication of the Federation of American Societies for Experimental Biology vol. 34,10 (2020): 13106–13124. doi:10.1096/fj.202000767R

50. van der Windt, Gerritje J W et al. “Mitochondrial respiratory capacity is a critical regulator of CD8+ T cell memory development.” Immunity vol. 36,1 (2012): 68–78. doi: 10.1016/j.immuni.2011.12.007

51. Desdín-Micó, Gabriela et al. “Mitochondrial activity in T cells.” Mitochondrion vol. 41 (2018): 51-57. doi: 10.1016/j.mito.2017.10.006

52. Pfleger, J., He, M. & Abdellatif, M. Mitochondrial complex II is a source of the reserve respiratory capacity that is regulated by metabolic sensors and promotes cell survival.Cell Death Dis 6, e1835 (2015).

53. Mudd, Joseph C, and Michael M Lederman. “CD8 T cell persistence in treated HIV infection.” Current opinion in HIV and AIDS vol. 9,5 (2014): 500–5. doi:10.1097/COH.0000000000000086

54. Konjar, Špela, et al. “Intestinal Barrier Interactions with Specialized CD8 T Cells. Frontiers in Immunology. vol. 8 1281. 11 Oct. 2017, doi:10.3389/fimmu.2017.01281

55. Sathaliyawala T, et al. Distribution and compartmentalization of human circulating and tissue-resident memory T cell subsets. Immunity. 2013; 38:187–197.

56. Thome JJ, et al. Spatial map of human T cell compartmentalization and maintenance over decades of life. Cell. 2014; 159:814–828.

57. Coetzee, Lelani et al. “HIV, antiretroviral therapy and non-communicable diseases in sub-Saharan Africa: empirical evidence from 44 countries over the period 2000 to 2016.” Journal of the International AIDS Society vol. 22,7 (2019): e25364. doi:10.1002/jia2.25364

58. Lerner, Andrea M et al. “Comorbidities in Persons With HIV: The Lingering Challenge.” JAMA vol. 323,1 (2020): 19–20. doi:10.1001/jama.2019.19775

59. Kumari, Ruchi, and Parmjit Jat. “Mechanisms of Cellular Senescence: Cell Cycle Arrest and Senescence Associated Secretory Phenotype.” Frontiers in cell and developmental biology vol. 9 645593. 29 Mar. 2021, doi:10.3389/fcell.2021.645593

60. Desdín-Micó, Gabriela et al. “T cells with dysfunctional mitochondria induce multimorbidity and premature senescence.” Science (New York, N.Y.) vol. 368,6497 (2020): 1371–1376. doi:10.1126/science.aax0860

61. Soto-Heredero, Gonzalo et al. “Mitochondrial dysfunction defines T cell exhaustion.” Cell metabolism vol. 33,3 (2021): 470–472. doi: 10.1016/j.cmet.2021.02.010

62. Perl, Andras et al. “Mitochondrial hyperpolarization: a checkpoint of T-cell life, death and autoimmunity.” Trends in immunology vol. 25,7 (2004): 360–7. doi:10.1016/j.it.2004.05.001

63. Trapecar, Martin et al. “An Optimized and Validated Method for Isolation and Characterization of Lymphocytes from HIV+ Human Gut Biopsies.” AIDS research and human retroviruses vol. 33, S1 (2017): S31–S39. doi:10.1089/AID.2017.0208

64. Shacklett BL, Critchfield JW, Lemongello D. Isolating mucosal lymphocytes from biopsy tissue for cellular immunology assays. Methods Mol Biol. 2009; 485:347– 356.

65. Shacklett BL, et al. Optimization of methods to assess human mucosal T-cell responses to HIV infection. J Immunol Methods. 2003; 279:17–31.

66. Asowata, Osaretin E et al. “Irreversible depletion of intestinal CD4+ T cells is associated with T cell activation during chronic HIV infection. JCI insight. vol. 6,22 e146162. 22 Nov. 2021, doi: 10.1172/jci.insight.146162

67. Love, M.I., Huber, W. & Anders, S. Moderated estimation of fold change and dispersion for RNA-seq data with DESeq2. Genome Biol 15, 550 (2014).

68. Subramanian, Aravind et al. “Gene set enrichment analysis: a knowledge-based approach for interpreting genome-wide expression profiles.” Proceedings of the National Academy of Sciences of the United States of America vol. 102,43 (2005): 15545–50.

