## Supplemental figures for "Colon resident CD8+ T cells from people with HIV on antiretroviral therapy exhibit mitochondrial dysfunction and impaired responses"

Supplementary Figure 1

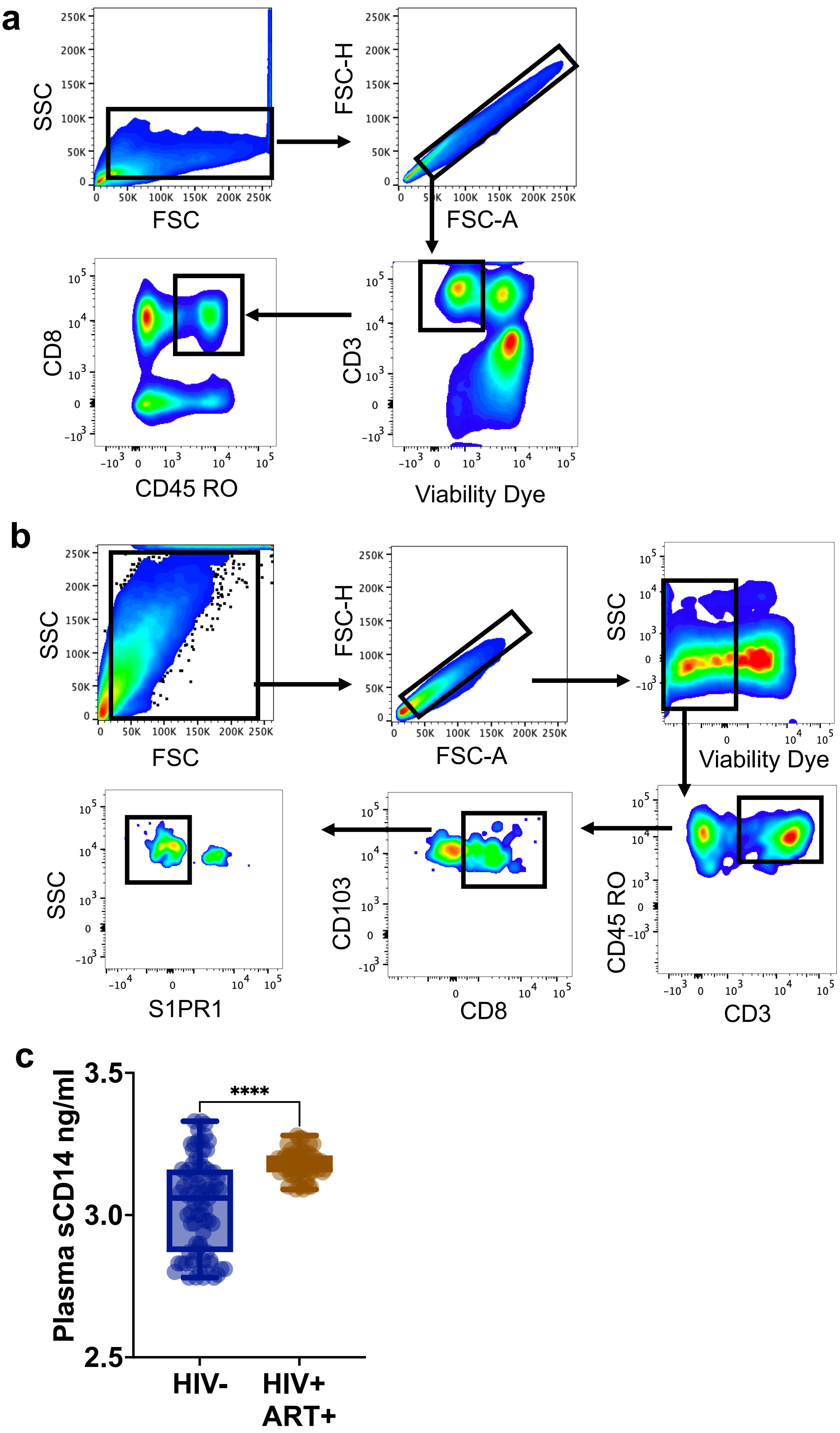

### Supplementary Figure 2

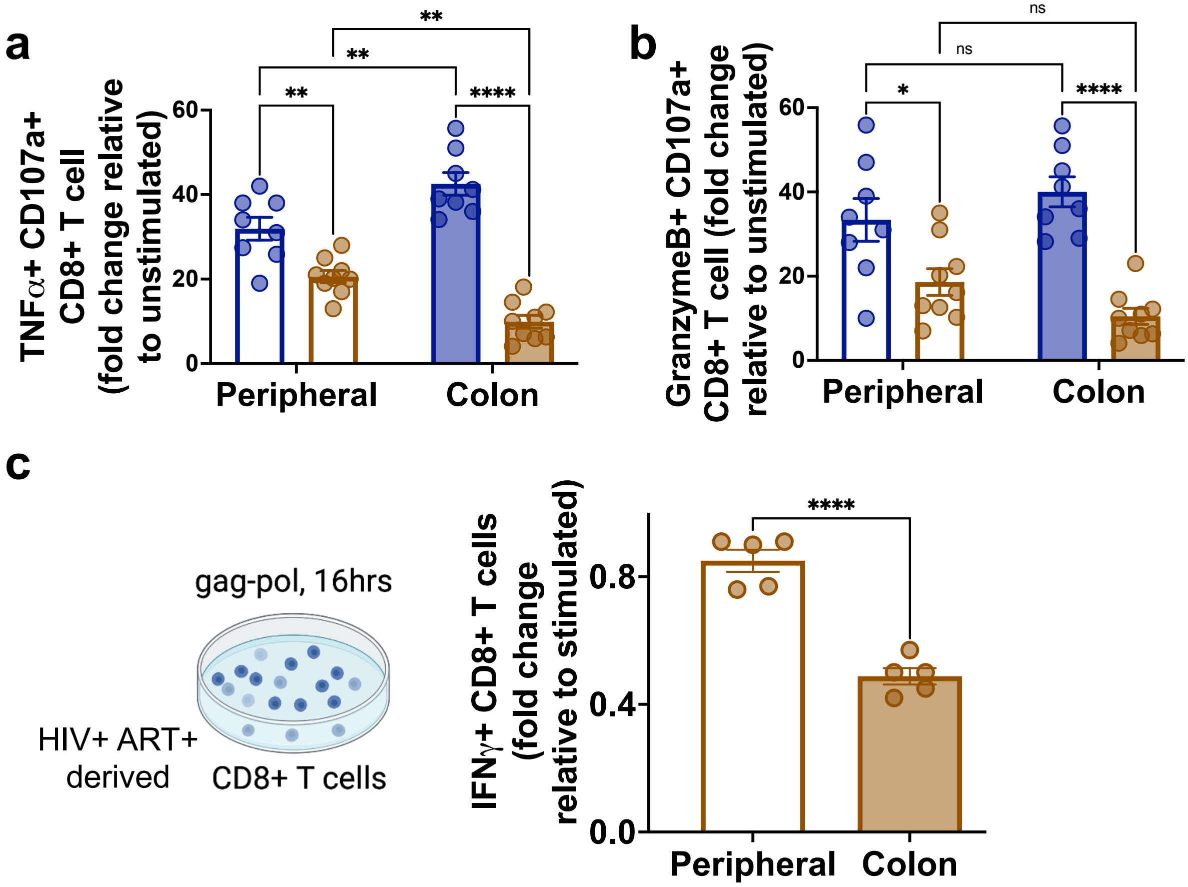

### Supplementary Figure 3

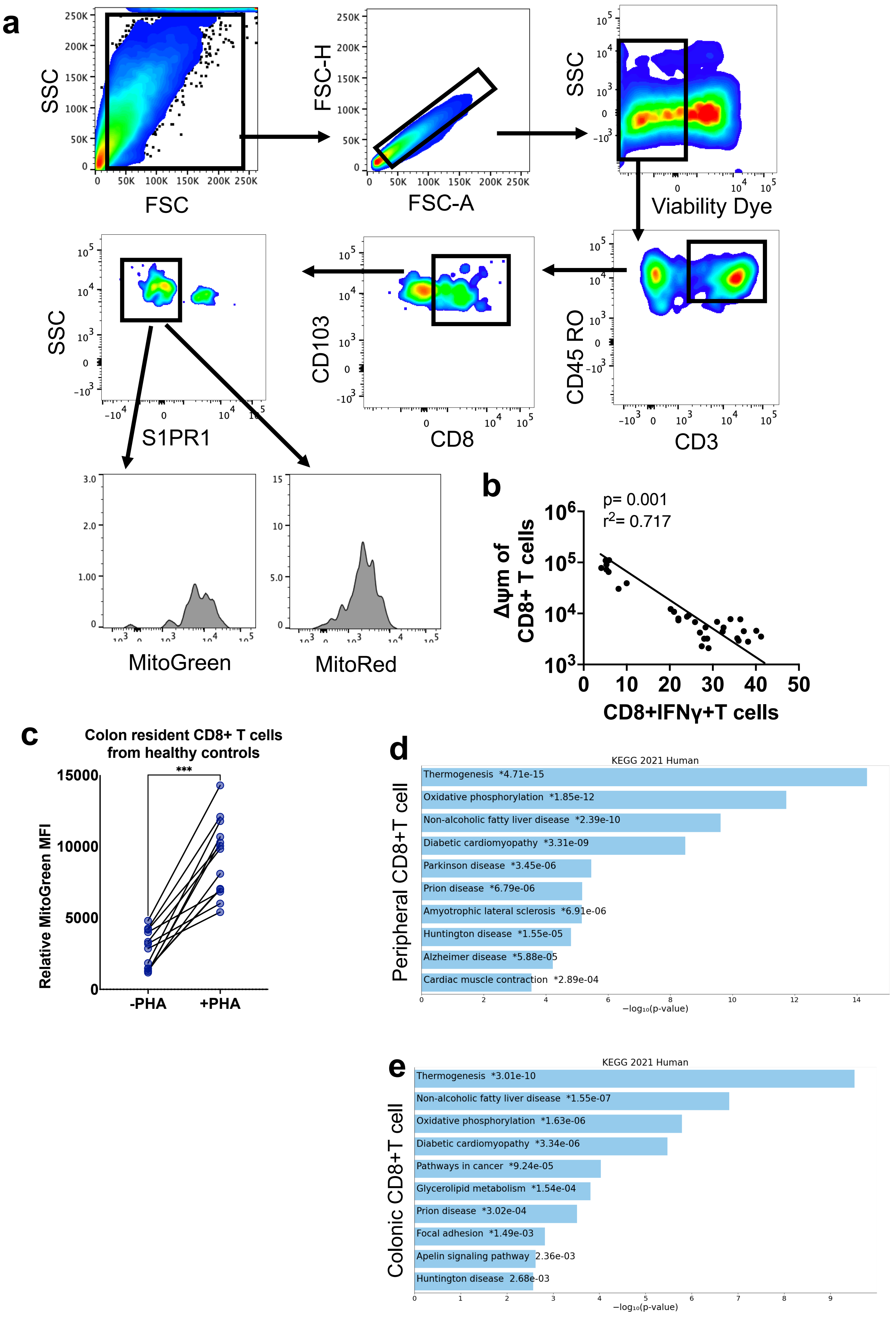

### Supplementary Figure 4

**a**

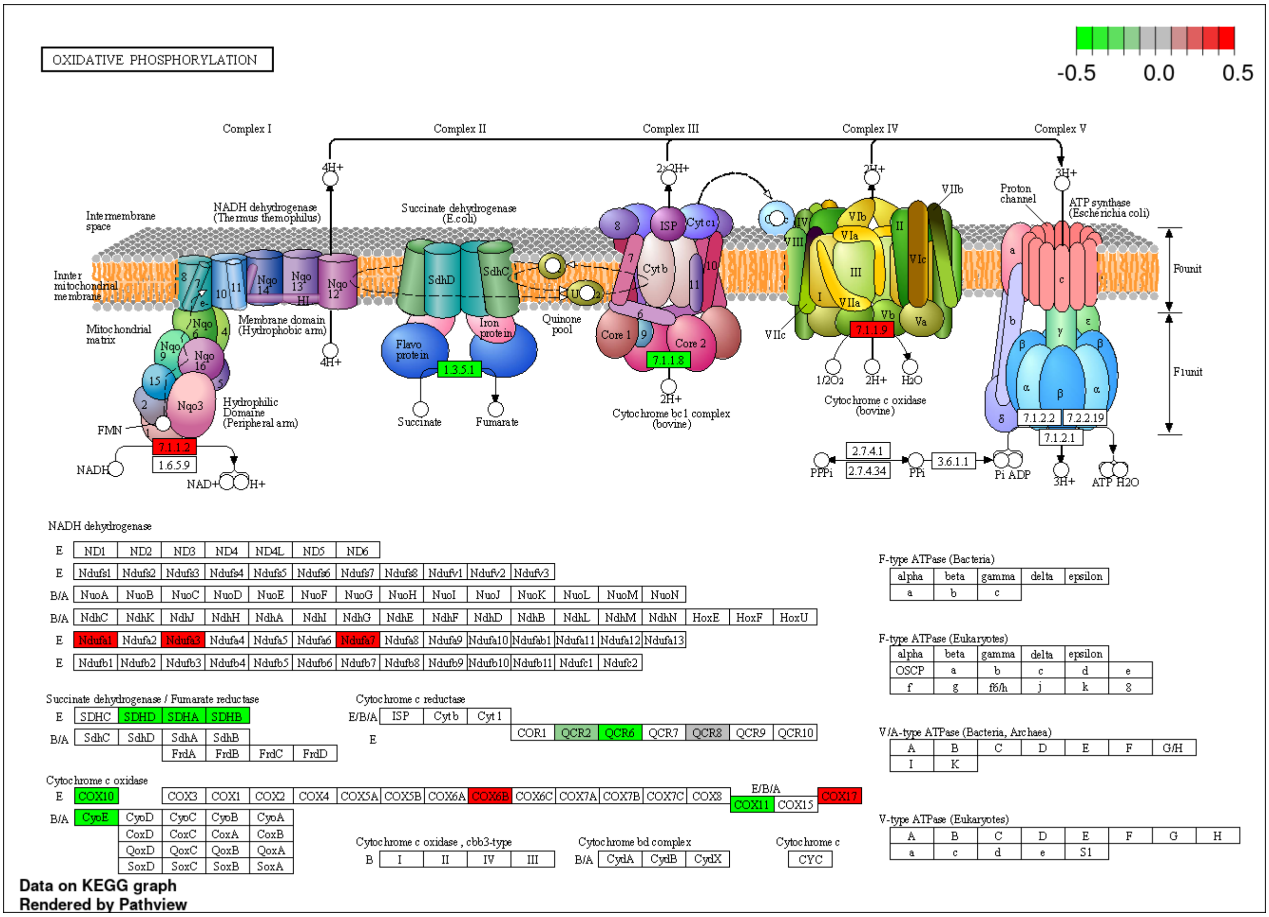


**b**

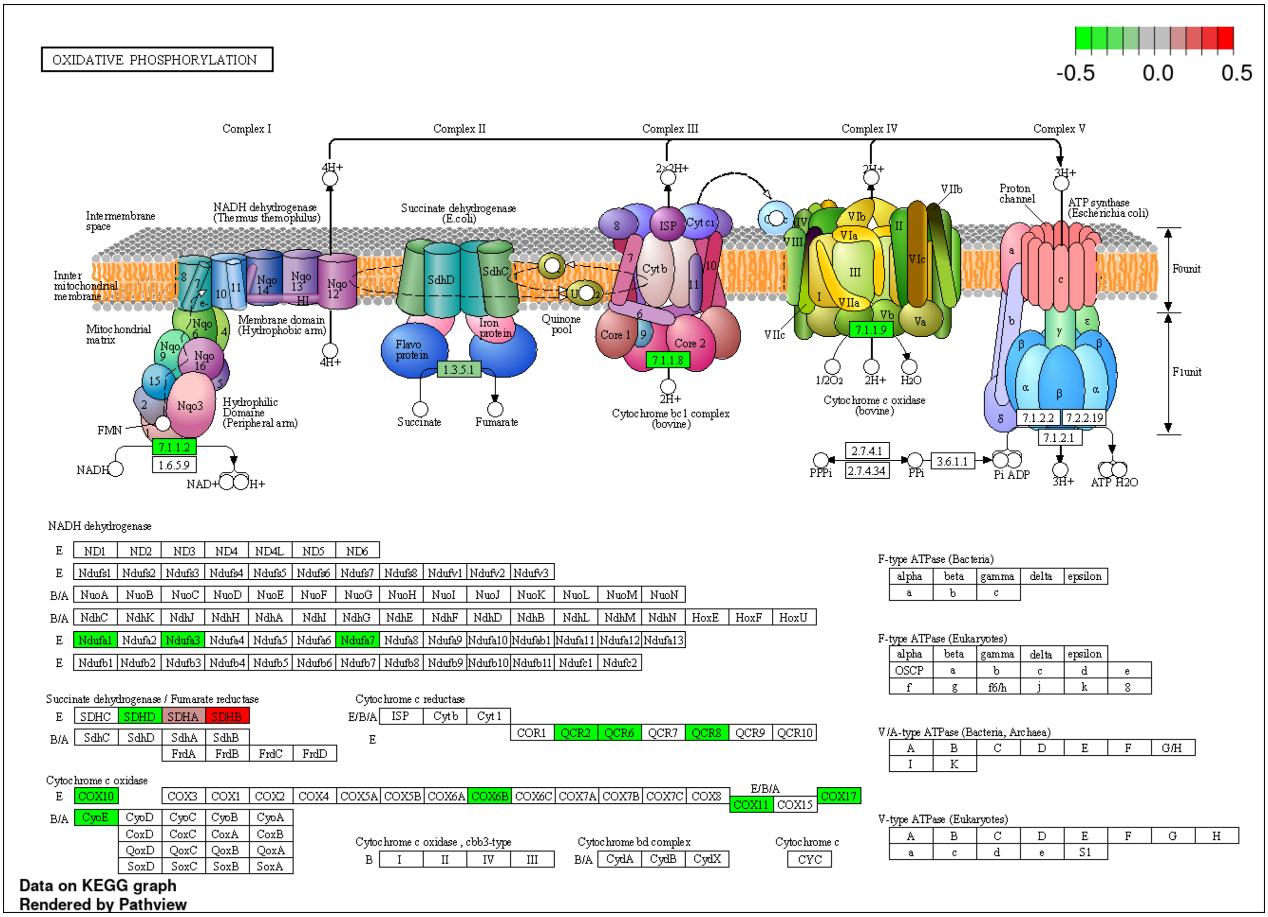

**Supplementary Figures**

Figure S1 related to Figure 1

a. Schematic representation of flow cytometric gating of peripheral CD8+ T cells
b. Schematic representation of flow cytometric gating of colonic CD8+ T cells
c. Plasma sCD14 measurement of HIV uninfected healthy donors and PWH on ART

Figure S2 related to Figure 2

a. Quantification of peripheral and colon resident CD8+ T (CD45RO+) cell in HIV uninfected healthy donors and PWH on ART for TNFa+/CD107a+ populations.
b. Quantification of peripheral and colon resident CD8+ T (CD45RO+) cell in HIV uninfected healthy donors and PWH on ART for granzymeB+/CD107a+ populations.
c. Schematic of ELISPOT assay; Quantification of the IFNg producing gag-pol antigen specific CD8+T cell in peripheral and colon resident CD8+T cell in PWH on ART.

Figure S3 related to Figure 3

a. Schematic representation of flow cytometric gating of colonic CD8+ T cells showing florescence intensity of Mito Red and Mito Green
b. Correlation between mitochondrial hyperpolarization and IFNg producing colonic CD8+ T cells independent of HIV status.
c. Quantification of MitoGreen MFI of healthy donor derived colonic CD8+ T cells with and out ex vivo stimulation by PHA
d. Bar chart of top enriched signaling pathway expressed transcriptionally in peripheral CD8+ T cells using the KEGG_Human gene set library
e. Bar chart of top enriched signaling pathway expressed transcriptionally in colonic CD8+ T cells using the KEGG_Human gene set library

Figure S4 Representation of mitochondrial electron transport chain (ETC) demonstrating the differential gene expression related to ETC complexes

a. between colonic CD8+ T cells from HIV uninfected healthy donors and PWH on ART
b. between peripheral CD8+ T cells from HIV uninfected healthy donors and PWH on ART
